# The genome sequence of Pavonine herpesvirus 1, a novel *Mardivirus* infecting peafowl

**DOI:** 10.64898/2026.02.16.706139

**Authors:** Steven R. Fiddaman, Soraia Barbosa, Miguel Carneiro, Joel M. Alves, Adrian L. Smith

## Abstract

Avian alphaherpesviruses of the genus *Mardivirus* include several economically important pathogens, such as the tumour-causing Marek’s disease virus (MDV-1) of chickens. However, the diversity of related viruses in wild and non-model avian hosts remains poorly characterised. Here, we describe the genome of Pavonine herpesvirus 1 (PaHv1), a novel *Mardivirus* infecting peafowl (*Pavo* spp.). Through de novo assembly of reads generated from whole-genome sequencing of 370 captive peafowl, we generated a single viral genomic scaffold of 134,149 bp in length and an estimated total genome size of ∼146 kbp following reconstruction of terminal repeat regions. PaHv1 exhibits canonical *Mardivirus* UL-IRL-IRS-US genomic architecture and encodes 169 open reading frames, of which 82 have clear orthologs in other *Mardivirus* species. Comparative genomic analyses revealed strong syntenic concordance between PaHv1, MDV-1, MDV-2 and Meleagrid alphaherpesvirus 1 (HVT), but with notable absences of the Meq oncogene, viral interleukin-8 (vIL8), and a truncated pp38 gene – genes known to underpin oncogenicity and virulence in MDV-1. Phylogenetic analysis places PaHv1 firmly within the *Mardivirus* genus as a sister to the poultry-associated MDV-1 and MDV-2 viruses, but distinct from a previously described Phasianid herpesvirus with a partial DNA terminase sequence. PaHv1 was detected predominantly in feather-derived samples, consistent with replication or persistence in the feather follicle epithelium. Taken together, these findings identify PaHv1 as a novel, likely non-oncogenic *Mardivirus*, expanding our understanding of herpesvirus diversity, evolution and host association in galliform birds.

**Importance:** Herpesviruses cause several economically important diseases in domesticated birds, including Marek’s disease, duck plague and infectious laryngotracheitis. Despite this, our understanding of herpesviruses infecting wild and semi-wild birds remains limited. Here, we present the genome of Pavonine herpesvirus 1 (PaHv1), a virus in the *Mardivirus* genus which infects peafowl. Based on comparative genomics with other *Mardivirus* species, we note the absence of several genes important for the high virulence of Marek’s Disease Virus, such as the Meq oncogene, viral interleukin-8 and a truncated pp38. This, taken together with absence of clinical disease in preliminary observations, suggests that PaHv1 causes either a subclinical infection or one with mild symptoms. Our work expands the known evolutionary breadth of avian herpesviruses in the *Mardivirus* genus. In addition, PaHv1 may provide a useful starting point for the development of novel vaccine vectors for pathogenic herpesvirus infections.

## Introduction

Herpesviruses comprise a family of large, enveloped, double-stranded DNA viruses that infect all major vertebrate lineages. A defining feature of these viruses is their ability to establish lifelong infection through latent reservoirs that can periodically reactivate to produce infectious virus and, in some cases, clinical disease. The family *Orthoherpesviridae* comprises three subfamilies: *Alphaherpesvirinae, Betaherpesvirinae* and *Gammaherpesvirinae*. Of these, the alphaherpesviruses are typically characterised by a short replication cycle, rapid destruction of infected cells, efficient spread within the host, and latency associated with neuronal or lymphoid tissues (Widén et al. 2012). Reactivation of latent alphaherpesviruses is often linked to physiological stress and can result in overt clinical disease.

Most well characterised avian herpesviruses (AHV) are alphaherpesviruses, particularly in the *Iltovirus* and *Mardivirus* genera. AHV are widespread in wild and domesticated birds, and although many cause subclinical or latent infections, there are several notable exceptions which result in severe and varied disease presentations. These include Marek’s disease (caused by Gallid alphaherpesvirus 2, genus: *Mardivirus*), which results in lymphoid tumours and paralysis in chickens (Nair 2018), duck plague (caused by Anatid alphaherpesvirus 1, genus: *Mardivirus*), a high-mortality disease with relatively non-specific clinical signs (Dhama et al. 2017), and infectious laryngotracheitis (caused by Gallid alphaherpesvirus 1, genus: *Iltovirus*), which causes predominantly respiratory symptoms in a variety of galliform hosts (Mo and Mo 2025). Disease outcomes in many avian herpesvirus infections are determined by the host species, the viral strain, the health of the host and environmental conditions.

In addition to the economically important AHV infecting domesticated poultry, a number of related viral species have been found in wild birds. These include Columbid herpesvirus 1 (pigeon herpesvirus), Acciptrid herpesvirus 1 (bald eagle herpesvirus), Ciconiid herpesvirus 1 (black stork herpesvirus), Falconid herpesvirus (causative agent of falcon inclusion body disease) and Psittacid herpesvirus 1 (Pacheco’s parrot disease virus) (Kaleta and Docherty 2008). A further putative species of *Mardivirus*, Phasianid herpesvirus 1, has been suggested to cause hepatocellular necrosis and mortality in at least three species of pheasants: mountain peacock pheasant (*Polyplectron inopinatum*), Malayan peacock pheasant (*Polyplectron malacense*) and Congo peafowl (*Afropavo congensis*) (Seimon et al. 2012), although the full genome of this virus has yet to be elucidated. Alphaherpesviruses are therefore widely distributed across avian orders, but remain unevenly characterised outside domesticated hosts.

To further improve our understanding of AHV in wild birds, we herein describe the genome of the tentatively named *Pavonine herpesvirus 1* (PaHv1), a *Mardivirus* species which infects peafowl (*Pavo* spp.). The virus we describe is distinct from the previously described Phasianid herpesvirus 1 (Seimon et al. 2012), despite an overlapping host range.

## Materials and methods

### Samples and mapping

We used whole-genome Illumina raw data (150bp paired-end reads) generated in (Barbosa et al. 2026), consisting of blood samples or growing feathers collected from 370 captive-raised Indian and green peafowl (*Pavo* spp.) (PRJNA1216069). To identify samples containing evidence of *Mardivirus* DNA, reads were aligned to the Marek’s Disease Virus 1 (MDV-1) genome (Gallid alphaherpesvirus 2; strain: RB-1B; accession: EF523390.1) using BWA mem (v0.7.17-r1188) (Li and Durbin 2009) with default parameters. Duplicate reads were flagged for downstream analysis using the *MarkDuplicates* function in Picard (v3.0.0) (http://broadinstitute.github.io/picard). Read group information was added to each BAM file using the Picard function *AddOrReplaceReadGroups*. To distinguish between samples with true MDV-1 infection and those with a distinct *Mardivirus*, we excluded any samples with reads mapping with an average per-read identity of >90% to MDV-1 from the subsequent de novo assembly. Viral-positive samples were then mapped to the peafowl genome and unmapped reads taken forward for de novo assembly.

### De novo assembly of Pavonine Herpesvirus 1 genome

Reads not mapping to the peafowl genome were extracted from BAM files using bedtools bamtofastq v2.26.0 (Quinlan and Hall 2010). Pooled reads from herpesvirus-positive (MDV-1 excluded) samples were assembled into a de novo assembly using SPAdes (v3.13.0) with read correction (Prjibelski et al. 2020) using the following parameters: -k 21,33,55,77,99,127 --careful --cov-cutoff off. Each assembled contig was then compared against the genomes for peafowl (GCA_045791835.1), human (GRCh38 assembly), Gallid alphaherpesvirus 2 (strain: RB-1B; accession: EF523390.1), Gallid alphaherpesvirus 3 (strain: HPRS24; accession: AB049735.1) and Meleagrid alphaherpesvirus 1 (strain: FC126 (Burmester); accession: NC_002641.1) using tBLASTx. Contigs with greatest identity to one of the three Mardivirus genomes and no significant match to the human or peafowl genomes were retained for further analysis. We further confirmed viral contigs using Prodigal (Hyatt et al. 2010) to identify open reading frames, followed by Diamond BLAST (Buchfink et al. 2015) using all RefSeq viral proteins as a reference database. All putative Mardivirus contigs were aligned to the genomes of Gallid alphaherpesvirus 2, Gallid alphaherpesvirus 3 and Meleagrid alphaherpesvirus 1 to identify the likely contig position. Contigs were then manually assembled using the other Mardivirus genomes as scaffolds.

### Refinement of viral genome assembly

Once a scaffold of the Pavonine herpesvirus genome had been created, the assembly went through a process of iterative refinement. FASTQ reads were mapped against the scaffold using BWA mem (Li and Durbin 2009), followed by sorting and indexing of the resulting BAM file using Samtools v1.15.1 (Li et al. 2009). At each of four stages of iterative refinement, junctions between contigs and genome termini were updated based on the consensus of mapped reads.

### Quality and completeness assessment

We evaluated the completeness and structural integrity of the assembled Pavonine herpesvirus genome using CheckV v1.0.1 (Nayfach et al. 2021). For the purposes of completeness estimates, we manually reconstructed putative terminal repeat regions (terminal repeat long, TRL; terminal repeat short, TRS), which are inverted copies of internal repeat long (IRL) and internal repeat short (IRS). The genome sequence, including manually curated terminal repeat regions consistent with herpesvirus genome architecture, was analysed in *end-to-end* mode against the CheckV reference database. CheckV estimated completeness and possible host contamination using a combination of amino acid identity (AAI)-based expected genome length, viral hallmark gene detection, and structural repeat analysis.

### Gene identification and annotation

Using Geneious Prime v2025.1.3, we identified all open reading frames (ORFs) > 240 bp (80 amino acids) in length in the Pavonine herpesvirus genomic sequence. All putative ORFs were then analysed in a custom Python script to determine likely orthologous genes. Translated ORF sequences from Pavonine herpesvirus, Meleagrid alphaherpesvirus 1, Gallid alphaherpesvirus 2 and Gallid alphaherpesvirus 3 were reciprocally compared against each other using BLASTp with an E-value threshold of 1e-5. Reciprocally identified ORFs were designated as orthologous genes. All Pavonine herpesvirus ORFs with at least one reciprocated hit in other viral species were retained for annotation. Finally, percentage identity at the amino acid level was calculated for each orthologous gene. Pavonine herpesvirus genes with clear orthologs in other species were named according to established herpesvirus gene nomenclature.

### Phylogenetic analysis

We identified six genes in the unique long (UL) region of the Pavonine herpesvirus genome which are present as single-copy orthologs in all known herpesvirus genomes – UL15, UL19, UL22, UL27, UL28, UL30. Coding sequences from these genes were extracted and translated from seven other herpesviruses: Herpes simplex virus 1 (HSV1; OQ724913.1), Psittacid herpesvirus 1 (NC_005264), Columbid alphaherpesvirus 1 (KX589235), Falconid herpesvirus 1 (KJ668231.1), Meleagrid herpesvirus 1 (NC_002641), Gallid alphaherpesvirus 2 (EF523390.1), and Gallid alphaherpesvirus 3 (HQ840738). Protein alignments were generated for each of these loci using Muscle v5.1 (Edgar 2004), then concatenated to generate a final alignment of 6592 amino acids. A maximum likelihood tree was then generated using RAxML v8.2.11 (Stamatakis 2014) using the GAMMA BLOSUM62 protein model and 100 bootstrap replicates.

To compare Pavonine herpesvirus 1 to a previously reported Phasianid herpesvirus affecting mountain peacock pheasant (*Polyplectron inopinatum*), Malayan peacock pheasant (*Polyplectron malacense*), and the Congo peafowl (*Afropavo congensis*), we aligned the partial (162 amino acid) DNA terminase sequence (UL15) reported in (Seimon et al. 2012) (accession: JN127369), to the seven herpesviruses listed above. Sequences were aligned using Muscle and a maximum likelihood tree was generated using RAxML as described above.

### Per-sample positivity

To determine which peafowl samples contained PaHv1 reads, we mapped reads from each sample against the final PaHv1 genome using BWA mem (Li and Durbin 2009). Samples were considered to be positive for PaHv1 if they contained at least five unique mapped reads with an average per-read percentage identity of >98% to the reference. Enumerated mapped reads and percentage identities can be found in **Supplementary Table S1**.

### Data availability

The assembled genomic sequence of Pavonine herpesvirus 1 has been deposited in Genbank (accession PX991849). Raw reads used to perform de novo assembly of PaHv1 have been deposited in SRA (accession PRJNA1216069).

## Results

### Assembly of Pavonine Herpesvirus 1

Following de novo assembly of reads and iterative refinement, we generated a single contiguous genomic scaffold for the putative Pavonine herpesvirus 1 (PaHv1) of 134,149 bp in length and GC content of 46.3% (**Fig. 1**). Since our assembly methodology relied on short reads (150bp paired-end), we were unable to resolve the terminal repeat regions (TR_L_ and TR_S_). However, by manually curating the putative terminal repeat regions from the internal repeat regions (IR_L_ and IR_S_), we estimate the true size of the PaHv1 genome to be approximately 146,430 bp. This is in line with, albeit slightly shorter than, other *Mardivirus* genomes: HVT (159 kbp), MDV-1 (178 kbp) and MDV-2 (166 kbp).

**Figure 1.**
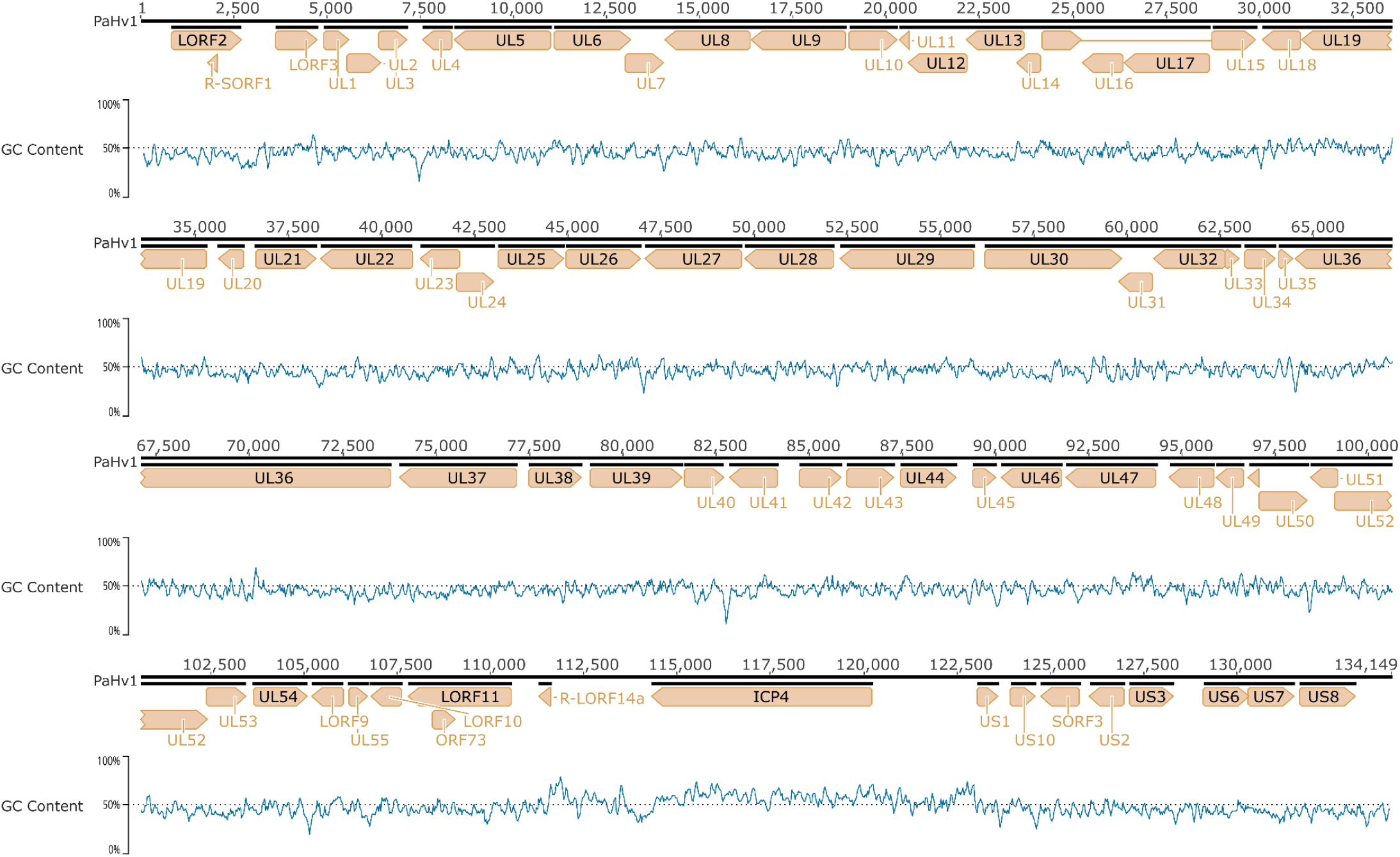
Genomic organisation of the PaHv1 genome. Open reading frames identified by homology are denoted by arrows pointing in the reading direction. GC content is indicated by the blue line.

CheckV classified the PaHv1 assembly as a complete, high-quality viral genome, with AAI- and HMM-based assessments supporting a completeness estimate of 100% with respect to gene content. CheckV also reported no evidence of contamination or non-viral open reading frames in the assembly. Mapping the original raw reads against the final assembly gave an average depth of coverage of 19.8× (±10.7). Taken together, these results support the high-confidence assembly of a structurally intact herpesvirus genome.

### Genomic organisation and Open Reading Frame (ORF) identification

The Pavonine herpesvirus 1 genome contained 169 putative ORFs of length >240 bp (80 amino acids). Based on reciprocal-best-hit BLAST analysis, we identified 82 ORFs with clear homology to an annotated ORF in at least one other *Mardivirus* genome (**Table 1**). Average percentage ORF identity was similar between PaHv1 and HVT, MDV-1 and MDV-2 (PaHv1 vs. HVT, 59.4%; PaHv1 vs. MDV-1, 64.0%; PaHv1 vs. MDV-2, 61.1%). In all three genomic comparisons, UL19 (major capsid protein) was the most highly conserved protein (average 83.9% amino acid identity). UL19 is a core herpesvirus capsid component and is highly conserved across Herpesviridae, forming the structural hexon/penton subunits of the icosahedral capsid (Bowman et al. 2003; Wang et al. 2018). The most divergent gene between PaHv1 and HVT/MDV-2 was envelope glycoprotein I (US7; average 38.4% amino acid identity), consistent with the general observation that gI/US7 is among the more rapidly evolving herpesvirus envelope proteins (Szpara et al. 2010).

**Table 1.** Orthology of open reading frames (ORFs) between Pavonine herpesvirus 1 and three other *Mardivirus* species: Herpesvirus of turkeys (HVT), Marek’s disease virus 1 (MDV-1) and Marek’s disease virus 2 (MDV-2). PaHv1 ORFs with at least one ortholog in at least one other species are listed. Length (in amino acids) and percentage identity are given for each ORF. ‘NA’ indicates that no ortholog was found for the ORF.

| PaHv1 | HVT (Meleagrid alphaherpesvirus 1) | MDV-1 (Gallid alphaherpesvirus 2) | MDV-2 (Gallid alphaherpesvirus 3) |
| --- | --- | --- | --- |
| LORF2; 621aa | HVT005 (lipase); 728aa; 48.4% | LORF2; 684aa; 57.0% | LORF1; 759aa; 52.1% |
| LORF3; 374aa | HVT007 (protein LORF2); 485aa; 48.0% | LORF3; 384aa; 39.9% | LORF2; 363aa; 40.3% |
| UL1; 229aa | HVT008 (envelope glycoprotein L); 197aa; 42.1% | UL1; 195aa; 59.3% | UL1; 190aa; 51.2% |
| UL2; 299aa | HVT009 (uracil-DNA glycosylase); 315aa; 63.3% | UL2; 313aa; 68.2% | UL2; 323aa; 63.1% |
| UL3; 256aa | HVT010 (nuclear protein UL3); 213aa; 63.5% | UL3; 228aa; 70.3% | UL3; 215aa; 70.8% |
| UL4; 262aa | HVT011 (nuclear protein UL4); 276aa; 57.0% | UL4; 268aa; 67.4% | UL4; 270aa; 62.0% |
| UL5; 859aa | HVT012 (helicase-primase helicase subunit); 858aa; 77.4% | UL5; 858aa; 83.1% | UL5; 857aa; 80.9% |
| UL6; 685aa | HVT013 (capsid portal protein); 729aa; 65.9% | UL6; 722aa; 69.2% | NA |
| UL7; 343aa | HVT014 (tegument protein UL7); 312aa; 60.3% | UL7; 305aa; 66.2% | UL7; 301aa; 62.5% |
| UL8; 761aa | HVT015 (helicase-primase subunit); 754aa; 58.4% | UL8; 769aa; 63.3% | UL8; 764aa; 63.1% |
| UL9; 842aa | HVT016 (DNA replication origin-binding helicase); 839aa; 70.5% | UL9; 841aa; 76.5% | UL9; 839aa; 71.0% |
| UL10; 424aa | HVT017 (envelope glycoprotein M); 423aa; 68.6% | UL10; 424aa; 79.7% | UL10; 424aa; 77.8% |
| UL11; 80aa | HVT018 (myristylated tegument protein); 81aa; 48.1% | UL11; 84aa; 50.0% | UL11; 81aa; 53.0% |
| UL12; 528aa | HVT019 (deoxyribonuclease); 521aa; 57.5% | UL12; 524aa; 56.2% | UL12; 525aa; 50.9% |
| UL13; 513aa | HVT020 (tegument serine/threonine protein kinase); 514aa; 74.4% | UL13A; 176aa; 49.2%<br>UL13B; 332aa; 77.3% | UL13; 500aa; 65.6% |
| UL14; 219aa | HVT021 (tegument protein UL14); 237aa; 51.4% | UL14; 243aa; 59.4% | UL14; 222aa; 55.1% |
| UL15; 741aa | HVT022 (DNA packaging terminase subunit 1); 738aa; 79.0% | UL15; 737aa; 81.4% | UL15; 748aa; 80.1% |
| UL16; 360aa | HVT023 (tegument protein UL16); 351aa; 62.0% | UL16; 360aa; 68.8% | UL16; 351aa; 66.9% |
| UL17; 762aa | HVT024 (DNA packaging tegument protein UL17); 728aa; 54.3% | UL17; 729aa; 59.7% | UL17; 722aa; 61.0% |
| UL18; 348aa | HVT025 (capsid triplex subunit 2); 320aa; 77.7% | UL18; 319aa; 82.1% | UL18; 321aa; 81.5% |
| UL19; 1392aa | HVT026 (major capsid protein); 1391aa; 82.7% | UL19; 1393aa; 85.9% | UL19; 1392aa; 83.2% |
| UL20; 232aa | HVT027 (envelope protein UL20); 230aa; 52.0% | UL20; 234aa; 63.7% | UL20; 234aa; 56.8% |
| UL21; 539aa | HVT028 (tegument protein UL21); 537aa; 49.6% | UL21; 546aa; 55.3% | UL21; 532aa; 56.1% |
| UL22; 814aa | HVT029 (envelope glycoprotein H); 808aa; 54.5% | UL22; 813aa; 60.7% | UL22; 838aa; 53.4% |
| UL24; 326aa | HVT030 (nuclear protein UL24); 292aa; 60.2% | UL24; 304aa; 51.8% | UL24; 316aa; 48.4% |
| UL23; 352aa | HVT031 (thymidine kinase); 350aa; 59.5% | UL23; 352aa; 73.7% | UL23; 352aa; 70.7% |
| UL25; 584aa | HVT032 (DNA packaging tegument protein UL25); 586aa; 66.3% | UL25; 583aa; 70.1% | UL25; 582aa; 72.3% |
| UL26; 661aa | HVT033 (capsid maturation protease); 643aa; 51.8% | UL26; 663aa; 57.3% | UL26; 639aa; 55.1% |
| UL27; 863aa | HVT035 (envelope glycoprotein B); 870aa; 79.6% | UL27; 865aa; 83.8% | UL27; 865aa; 80.3% |
| UL28; 793aa | HVT036 (DNA packaging terminase subunit 2); 787aa; 67.8% | UL28; 793aa; 73.9% | UL28; 787aa; 72.1% |
| UL29; 1192aa | HVT037 (single-stranded DNA-binding protein); 1190aa; 81.0% | UL29; 1191aa; 81.1% | UL29; 1190aa; 77.6% |
| UL30; 1218aa | HVT038 (DNA polymerase catalytic subunit); 1205aa; 72.0% | UL30; 1220aa; 73.7% | UL30; 1211aa; 72.9% |
| UL31; 305aa | HVT039 (nuclear egress lamina protein); 305aa; 74.8% | UL31; 300aa; 77.4% | UL31; 304aa; 76.2% |
| UL33; 135aa | HVT040 (DNA packaging protein UL33); 135aa; 60.7% | UL33; 134aa; 62.3% | UL33; 131aa; 63.1% |
| UL32; 633aa | HVT041 (DNA packaging protein UL32); 632aa; 68.0% | UL32; 641aa; 71.6% | UL32; 626aa; 68.4% |
| UL34; 279aa | HVT042 (nuclear egress membrane protein); 277aa; 69.4% | UL34; 277aa; 65.0% | UL34; 279aa; 63.3% |
| UL35; 130aa | HVT043 (small capsid protein); 130aa; 64.5% | UL35; 131aa; 71.4% | UL35; 127aa; 65.3% |
| UL36; 3099aa | HVT044 (large tegument protein); 3112aa; 52.6% | UL36; 3357aa; 61.3% | UL36; 3068aa; 60.3% |
| UL37; 1046aa | HVT045 (tegument protein UL37); 1041aa; 61.1% | UL37; 1046aa; 68.5% | UL37; 1046aa; 61.1% |
| UL38; 468aa | HVT046 (capsid triplex subunit 1); 470aa; 66.0% | UL38; 470aa; 66.8% | UL38; 470aa; 66.8% |
| UL39; 822aa | HVT047 (ribonucleotide reductase subunit 1); 820aa; 77.9% | UL39; 822aa; 81.5% | UL39; 797aa; 76.8% |
| UL40; 351aa | HVT048 (ribonucleotide reductase subunit 2); 355aa; 80.5% | UL40; 343aa; 76.0% | UL40; 353aa; 76.9% |
| UL41; 433aa | HVT049 (tegument host shutoff protein); 428aa; 65.3% | UL41; 441aa; 69.3% | UL41; 423aa; 64.9% |
| UL42; 369aa | HVT050 (DNA polymerase processivity subunit); 369aa; 70.1% | UL42; 369aa; 79.1% | UL42; 369aa; 71.3% |
| UL43; 423aa | HVT051 (envelope protein UL43); 422aa; 47.5% | UL43; 420aa; 50.0% | UL43; 414aa; 49.2% |
| UL44; 497aa | HVT052 (envelope glycoprotein C); 489aa; 73.2% | UL44B; 284aa; 84.6% | UL44; 478aa; 69.1% |
| UL45; 211aa | HVT053 (membrane protein UL45); 212aa; 70.8% | UL45; 211aa; 78.7% | UL45; 210aa; 65.4% |
| UL46; 535aa | HVT054 (tegument protein VP11/12); 494aa; 41.0% | UL46; 568aa; 40.8% | UL46; 581aa; 43.8% |
| UL47; 807aa | HVT055 (tegument protein VP13/14); 788aa; 45.2% | UL47; 808aa; 53.8% | UL47; 811aa; 49.5% |
| UL48; 393aa | HVT056 (transactivating tegument protein VP16); 422aa; 55.8% | UL48; 427aa; 63.7% | UL48; 424aa; 57.1% |
| UL49; 242aa | HVT057 (tegument protein VP22); 283aa; 51.6% | UL49; 249aa; 53.8% | UL49; 241aa; 54.5% |
| UL50; 435aa | HVT058 (deoxyuridine triphosphatase); 437aa; 60.2% | UL50; 436aa; 64.5% | UL50; 435aa; 59.3% |
| UL49.5; 96aa | HVT059 (envelope glycoprotein N); 94aa; 69.7% | UL49.5; 95aa; 62.5% | UL49.5; 95aa; 60.4% |
| UL52; 1101aa | HVT060 (helicase-primase primase subunit); 1078aa; 62.2% | UL52; 1074aa; 67.2% | UL52; 1071aa; 64.2% |
| UL51; 242aa | HVT061 (tegument protein UL51); 240aa; 58.6% | UL51; 249aa; 59.2% | UL51; 248aa; 62.4% |
| UL53; 353aa | HVT062 (envelope glycoprotein K); 349aa; 46.3% | UL53; 354aa; 64.4% | UL53; 354aa; 59.0% |
| UL54; 479aa | HVT063 (multifunctional expression regulator); 476aa; 49.1% | UL54; 473aa; 54.2% | UL54; 476aa; 48.8% |
| UL55; 168aa | HVT065 (nuclear protein UL55); 169aa; 55.9% | UL55; 166aa; 51.5% | UL55; 167aa; 57.1% |
| LORF10; 279aa | HVT066 (myristylated tegument protein CIRC); 185aa; 50.7% | LORF10; 194aa; 59.8% | NA |
| LORF9; 280aa | HVT067 (protein LORF4A); 265aa; 48.1% | LORF9; 269aa; 64.7% | LORF4; 268aa; 59.2% |
| LORF11; 920aa | HVT069 (protein LORF5); 883aa; 38.9% | LORF11; 903aa; 51.7% | LORF5; 873aa; 48.6% |
| R-LORF14a; 110aa | HVT071 (protein pp38); 84aa; 42.9% | R-LORF14a; 290aa; 44.2% | pp38; 249aa; 45.7% |
| US1; 190aa | HVT085 (regulatory protein ICP22); 165aa; 57.6% | US1; 179aa; 63.2% | US1; 173aa; 57.5% |
| US10; 221aa | HVT086 (virion protein US10); 209aa; 48.9% | US10; 213aa; 62.7% | US10; 231aa; 63.4% |
| SORF3; 349aa | HVT087 (protein SORF3); 346aa; 43.3% | SORF3; 351aa; 57.8% | SORF4; 322aa; 49.1% |
| US2; 306aa | HVT088 (virion protein US2); 282aa; 50.4% | US2; 270aa; 60.2% | US2; 271aa; 55.7% |
| US3; 388aa | HVT089 (serine/threonine protein kinase US3); 394aa; 60.0% | US3; 402aa; 61.6% | US3; 391aa; 55.9% |
| US6; 402aa | HVT090 (envelope glycoprotein D); 371aa; 48.4% | US6; 398aa; 55.8% | US6; 385aa; 50.5% |
| US7; 402aa | HVT091 (envelope glycoprotein I); 356aa; 37.9% | US7; 355aa; 48.2% | US7; 355aa; 38.9% |
| US8; 499aa | HVT092 (envelope glycoprotein E); 498aa; 44.4% | US8; 497aa; 48.5% | US8; 518aa; 42.4% |
| ICP4; 1964aa | HVT095 (transcriptional regulator ICP4); 2164aa; 43.0% | immediate early protein ICP4; 2321aa; 45.9% | ICP4; 2033aa; 52.9% |
| ORF_115; 116aa | NA | hypothetical protein; 60aa; 57.5% | NA |
| ORF_116; 71aa | NA | NA | ORF473 (hypothetical protein); 111aa; 50.8% |
| ORF_144; 69aa | NA | hypothetical protein; 128aa; 56.7% | NA |
| R-SORF; 141aa | NA | R-SORF1; 108aa; 60.9% | NA |
| ORF_73; 203aa | NA | hypothetical protein; 206aa; 49.8% | ORF361 (hypothetical protein); 162aa; 44.1% |
| ORF_105; 105aa | NA | hypothetical protein; 92aa; 58.9% | NA |
| ORF_126; 71aa | NA | LORF8; 208aa; 64.2% | NA |
| ORF_127; 79aa | NA | LORF8; 208aa; 67.1% | LORF8 (hypothetical protein LORF8); 62aa; 54.4% |
| ORF_62; 125aa | NA | LORF6; 155aa; 62.0% | NA |
| ORF_166; 79aa | NA | NA | ORF223 (hypothetical protein n025R); 92aa; 81.3% |
| ORF_176; 107aa | NA | NA | ORF273 (ORF273 protein); 273aa; 48.6% |

The gross genomic organisation of PaHv1 is in line with other *Mardivirus* species. While we were not able to explicitly resolve the terminal repeat regions (TR_L_ and TR_S_), we found that the viral genomic scaffold comprises a long unique (U_L_) region, long internal repeat (IR_L_), short internal repeat (IR_S_) and short unique (U_S_) region in the expected orientation. Furthermore, genes within these regions largely follow strong syntenic concordance with other *Mardivirus* species (**Fig. 2**). One notable exception to this is the absence of a block of 10 genes in the long repeat region in PaHv1 which is present in MDV-1. This block of genes is also absent in HVT (Kingham et al. 2001) and contains the *Meq* oncogene and the v-IL8 gene, which are responsible for tumourigenesis and recruitment of chicken immune cells, respectively, in MDV-1 (Parcells et al. 2001; Jones et al. 1992). In addition, two further genes (SORF1 and SORF2) are not found in PaHv1 or HVT (Kingham et al. 2001) but are present in both MDV-1 and MDV-2. Finally, PaHv1 also shows a truncated pp38 (R-LORF14a) gene which is 110 amino acids in length, compared to MDV-1 (290 amino acids) and MDV-2 (249 amino acids). HVT pp38 is also truncated (84 amino acids) and has been reported previously (Kingham et al. 2001). These findings indicate that while PaHv1 has broadly conserved genomic architecture that is consistent with *Mardivirus* species and wider herpesviruses, there are key differences – notably in the repeat regions – between PaHv1 and the two chicken *Mardivirus* species, MDV-1 and MDV-2. Specifically, the absence of the *Meq* oncogene and the truncation of pp38, two loci essential for MDV-1-induced lymphomagenesis, suggest that PaHv1 is unlikely to possess the oncogenic potential characteristic of MDV-1.

**Figure 2.**
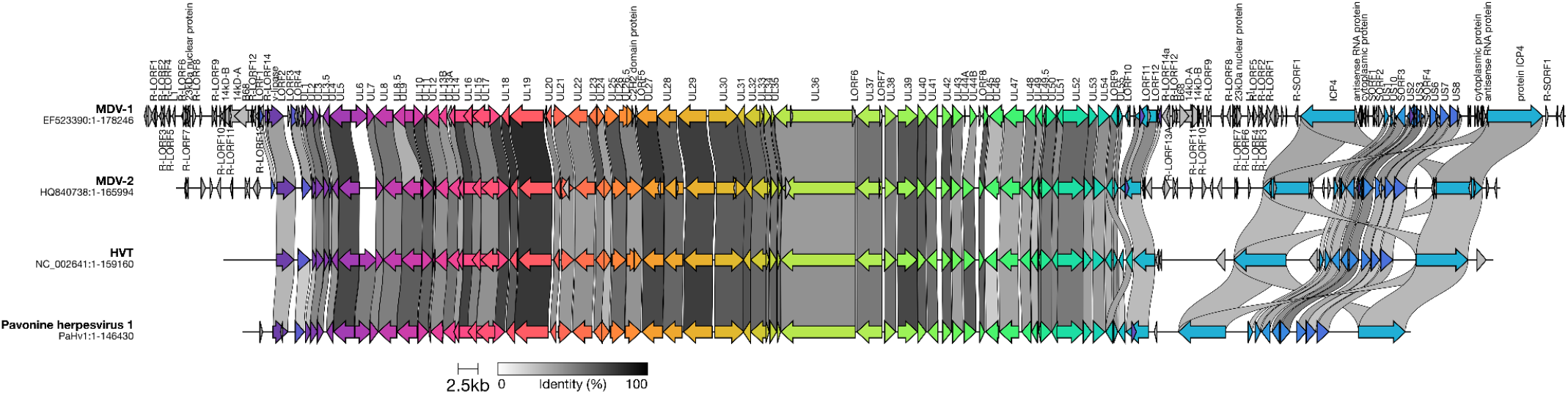
Synteny analysis of the PaHv1 genome with respect to other *Mardivirus* species (MDV-1, MDV-2 and HVT). Open reading frames (ORFs) are denoted by arrows. Coloured arrows are ORFs with an ortholog in at least one other species, linked by a grey band. The shade of grey band indicates the percentage identity between ORFs.

### Phylogenetic analysis of Mardivirus species

To confirm that PaHv1 sits within the *Mardivirus* genus, we constructed a phylogenetic tree of PaHv1 along with other *Mardivirus* and alphaherpesvirus outgroup species. The alignment used to build the phylogeny was based on concatenated amino acid alignments of core genes that are ubiquitous in herpesvirus species across different genera (UL15, UL19, UL22, UL27, UL28, UL30). The overall percentage identity between PaHv1 and HVT, MDV-1 and MDV-2 in the concatenated amino acid alignment was 71.4%, 75.7% and 72.3%, respectively. After rooting to Psittacid alphaherpesvirus 1 (genus: *Iltovirus*), the phylogeny of existing alphaherpesviruses followed previous observations (Verweij et al. 2015), with Herpes simplex virus 1 (genus: *Simplexvirus*) forming a sister group to Falconid herpesvirus, Columbid herpesvirus, and the rest of the *Mardivirus* genus (HVT, MDV-1 and MDV-2). Pavonine herpesvirus 1 lies within the *Mardivirus* clade as a sister to the chicken *Mardivirus* species, MDV-1 and MDV-2. The tree topology is very well supported (bootstrap values ≥ 96), confirming the position of PaHv1 as a member of the *Mardivirus* genus.

A previous report (Seimon et al. 2012) provides a partial sequence of the DNA terminase gene (UL15; accession: JN127369) from a putative herpesvirus infecting mountain peacock pheasant (*Polyplectron inopinatum*), Malayan peacock pheasant (*Polyplectron malacense*), and the Congo peafowl (*Afropavo congensis*), named Phasianid herpesvirus. To determine whether the Pavonine herpesvirus reported herein is the same as the previously published Phasianid herpesvirus, we aligned the DNA terminase fragment (162 amino acids) from Phasianid herpesvirus with the homologous region from PaHv1 and other alphaherpesviruses. Although the short alignment length meant support values in the resulting phylogeny were generally poor, Phasianid herpesvirus and PaHv1 were clearly different (**Fig. 3**) – Phasianid herpesvirus clusters closely with MDV-2 (93.8% amino acid identity) but had only 87.7% amino acid identity with PaHv1. Although the extent to which the host ranges of Phasianid herpesvirus and PaHv1 overlap remains unclear, the two viruses are clearly distinct, despite infecting genetically very similar hosts.

**Figure 3.**
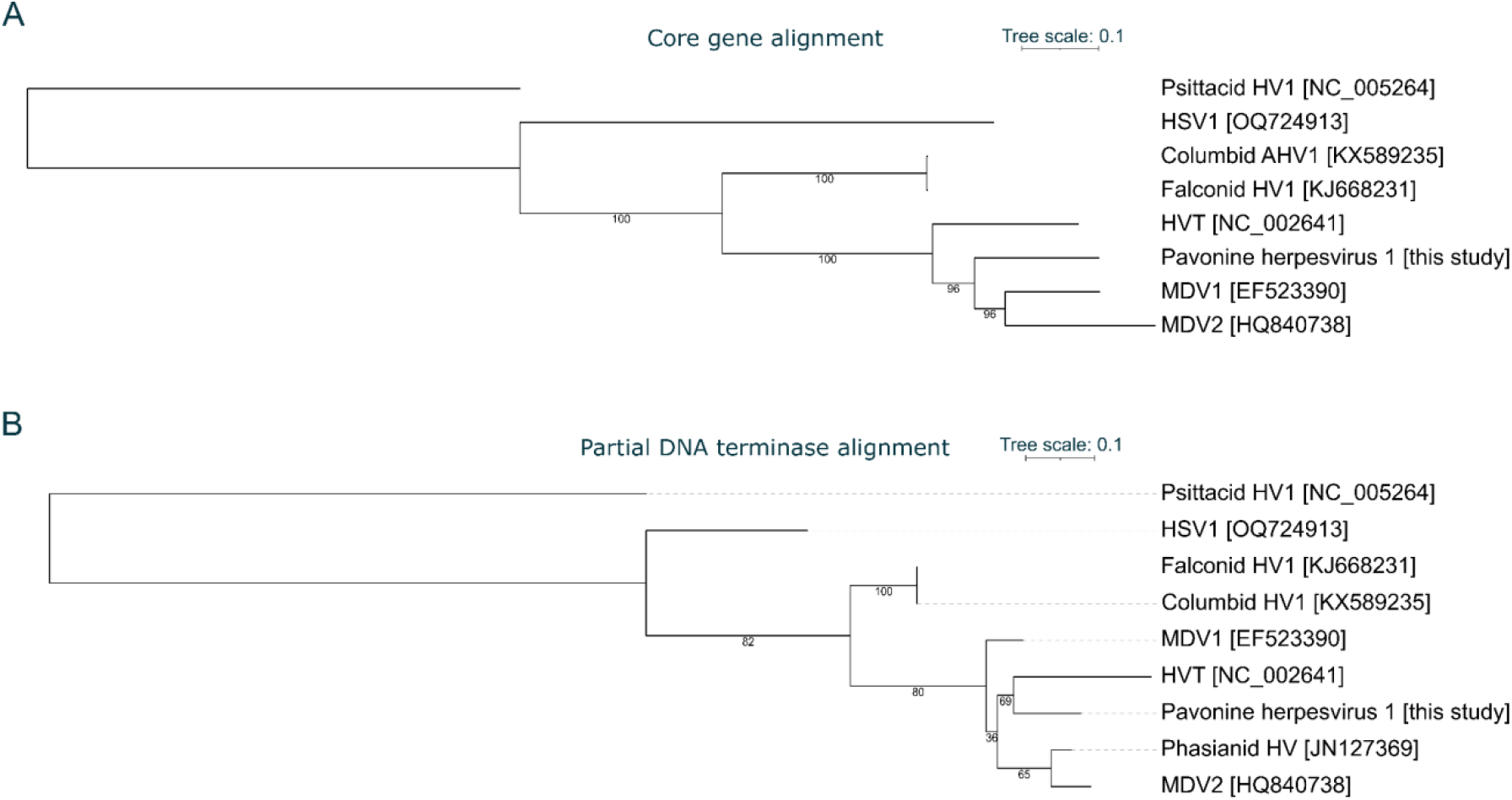
Phylogenetic analysis of Pavonine herpesvirus 1 and related viruses. (A) Maximum likelihood tree based on a concatenated amino acid alignment of six core genes (UL15, UL19, UL22, UL27, UL28, UL30). (B) Maximum likelihood tree based on an amino acid alignment of a partial DNA terminase sequence given in (Seimon et al. 2012); accession: JN127369) with eight other overlapping sequences from other herpesviruses. For both trees, branch labels are bootstrap support values and both trees were rooted to the Psattacid herpesvirus 1 sequence (subfamily: *Alphaherpesvirinae*; genus: *Iltovirus*). Abbreviations: HSV, Herpes simplex virus; MDV, Marek’s Disease Virus; HVT, Herpesvirus of Turkeys.

### Case positivity of Pavonine herpesvirus 1

We examined the characteristics of samples that were positive for PaHv1. Per-sample positivity was defined based on ≥5 independent reads mapping to the PaHv1 genome with an average percentage identity of >98%. Of the 370 peafowl with shotgun sequencing data, 27 were positive for PaHv1, giving an overall case positivity rate of 7.3% (**Table 2**). However, PaHv1 was substantially more common in feather samples than blood samples. While only 4/278 blood samples were positive for PaHv1 (1.4%), 23/92 feather samples (25%) were positive. These data indicate that, like MDV-1, PaHv1 is present at relatively high levels in feather-derived material, consistent with active replication or persistence in feather follicle epithelium.

**Table 2.** Case positivity of PaHv1 in peafowl samples (blood and feathers), defined as samples with ≥5 mapped reads and an average identity of >98% to the PaHv1 genome.

| Extraction tissue | Samples | PaHV1 | PaHv1 positivity (%) |
| --- | --- | --- | --- |
| Blood | 278 | 4 | 1.4 |
| Feathers | 92 | 23 | 25.0 |
| **Totals** | 370 | 27 | 7.3 |

## Discussion

We report the genomic sequence of the tentatively named Pavonine herpesvirus 1 (PaHv1), a novel member of the *Mardivirus* genus of alphaherpesviruses. PaHv1 has an estimated genome size of approximately 146 kbp and a canonical UL-IRL-IRS-US genomic architecture, consistent with other *Mardivirus* species and alphaherpesviruses (Dotto-Maurel et al. 2025). Comparative genomic and phylogenetic analyses clearly distinguish PaHv1 from all previously described Galliform herpesviruses, including Gallid alphaherpesvirus 2 (MDV-1), Gallid alphaherpesvirus 3 (MDV-2) and Meleagrid alphaherpesvirus 1 (HVT), as well as from the previously reported Phasianid herpesvirus with a partial DNA terminase fragment (Seimon et al. 2012).

Most notably among the *Mardivirus* species, MDV-1 has caused tumours in chickens from the 1920s, having previously only caused a comparatively mild paralytic syndrome (Fiddaman et al. 2023; Nair 2005). Frequently evolving to evade vaccine protection, MDV-1 causes substantial losses to the poultry industry (estimated at between US$1-2bn per year). A defining feature of PaHv1 is the absence of several genes that underpin the virulence and oncogenicity of MDV-1. Most notably, PaHv1 lacks the *Meq* oncogene, which encodes a basic leucine zipper transcription factor that is both necessary and sufficient for MDV-1-induced T-cell lymphomagenesis (Brown et al. 2009; Lupiani et al. 2004). Deletion or truncation of *Meq* in MDV-1 results in complete attenuation and loss of tumour formation *in vivo*, highlighting the essential role of this gene in oncogenesis. *Meq* is similarly absent from the other non-oncogenic Mardiviruses, HVT and MDV-2, and it is highly unlikely that PaHv1 is capable of inducing MDV-1-like lymphoid tumours.

In addition to the absence of *Meq*, PaHv1 contains a truncated version of *pp38* (R-LORF14a). In MDV-1, pp38 is expressed during both lytic infection and latency, and plays a critical role in maintaining transformed T cells and sustaining tumour growth (Chen et al. 1992). Functional studies have shown that pp38-null or truncated MDV-1 mutants exhibit reduced pathogenicity and impaired lymphoproliferation (Gimeno et al. 2005). Pp38 is similarly truncated in the non-oncogenic HVT, further underscoring the contribution of key genes to the development of lymphoid tumours in *Mardivirus* species.

Furthermore, PaHv1 lacks a homolog of viral interleukin-8 (*vIL8*), a chemokine encoded by MDV-1 that is crucial for recruiting activated CD4^+^ T cells to sites of infection, thereby facilitating viral spread and tumour initiation (Parcells et al. 2001). Experimental deletion of *vIL8* from MDV-1 significantly reduces early cytolytic infection and attenuates disease severity (Parcells et al. 2001).

The clear genetic distinction between PaHv1 and the previously reported Phasianid herpesvirus is particularly noteworthy given their potential overlap in host ranges. The clustering of the Phasianid herpesvirus UL15 fragment with MDV-2 rather than with PaHv1 suggests that multiple, deeply divergent *Mardivirus* lineages circulate among Phasianidae and related Galliform birds. This pattern is consistent with long-term virus-host co-divergence punctuated by occasional host switches, a mode of evolution that has been proposed for alphaherpesviruses more broadly (Brito et al. 2021). The relatively even amino acid divergence between PaHv1 and the three poultry-associated Mardiviruses further argues against a recent spillover from domestic birds, instead supporting the existence of a distinct peafowl-associated viral lineage.

These findings have implications for both herpesvirus evolution and disease ecology. The modular gain and loss of genes within the long and short repeat regions, particularly those involved in immune modulation and oncogenesis, appears to be a major driver of phenotypic diversity within the *Mardivirus* genus (Afonso et al. 2001). At the same time, the frequent co-housing of peafowl with chickens and other galliform species in captive or semi-domesticated settings raises the possibility of cross-species transmission or recombination, processes that are well documented among herpesviruses (Casto et al. 2020). While there is currently no evidence that PaHv1 poses a risk to domestic poultry, its discovery highlights the importance of characterising herpesvirus diversity in non-model avian hosts.

A limitation of this study is that PaHv1 was identified incidentally from whole-genome sequencing of peafowl, precluding a systematic assessment of clinical signs associated with infection. The previously reported Phasianid herpesvirus (Seimon et al. 2012) caused severe symptoms, including hepatocellular necrosis and death. No unexplained deaths were reported in the peafowl sequenced for this study, so it is likely that the clinical signs of PaHv1 infection, if any, are milder than Phasianid herpesvirus infection. One putative clinical sign anecdotally reported by the owner of birds in which PaHv1 was prevalent was blindness or visual impairment. However, other viral diseases of peafowl, such as infectious laryngotracheitis (ILT; caused by Gallid alphaherpesvirus 1) are also associated with ocular symptoms, such as conjunctivitis and ocular discharge (Widén et al. 2012; Gowthaman et al. 2020). It remains to be established whether blindness is a true symptom of PaHv1 infection, or whether this was an incidental finding of other infectious or non-infectious aetiology.

If PaHv1 is found to cause mild symptoms or an asymptomatic infection, this would be consistent with the broader biology of herpesviruses, which are characterised by lifelong latency, frequent subclinical infection, and disease manifestation that is often contingent on host immune status, co-infections, or environmental stressors (Seeber et al. 2018). In Galliform hosts, both HVT and MDV-2 are widely prevalent yet typically asymptomatic (or cause mild symptoms), despite efficient replication and transmission (Afonso et al. 2001; Neerukonda et al. 2019). It therefore remains plausible that PaHv1 represents a similarly persistent but largely benign virus of peafowl, with any observed pathology arising independently or requiring specific permissive conditions.

Since *Mardivirus* species can also confer cross-species disease protection when used as vaccines, PaHv1 may represent a potential starting point for the development of alternative vaccine vectors. PaHv1 has a slightly higher average percentage identity to MDV-1 (75.7%) than HVT does to MDV-1 (73.5%), and only slightly less than MDV-2 to MDV-1 (76.0%). Since both HVT and MDV-2 are used in vaccine formulations against MDV-1 (Reddy et al. 2017), it is plausible that PaHv1 could, in principle, form the basis of a vaccine platform against MDV-1. Novel vaccine candidates against MDV-1 are especially valuable since no vaccine to date has provided complete sterile immunity against infection (Reddy et al. 2017), driving evolution of the virus towards greater levels of virulence (Read et al. 2015).

In conclusion, PaHv1 represents a novel, likely non-oncogenic *Mardivirus* that expands the known evolutionary breadth of the genus. Its genomic features, particularly the absence of *Meq*, truncation of *pp38*, and lack of *vIL8*, strongly suggest a pathogenic profile more similar to HVT than to MDV-1. Future studies integrating serological surveys, experimental infection models, and transcriptomic analyses of lytic and latent infection will be essential to define the host range, transmission dynamics, and clinical relevance of PaHv1, and to further elucidate how variation in repeat-region gene content shapes virulence across the *Mardivirus* lineage. More generally, our work highlights the power of using shotgun sequencing data to identify and characterise pathogens – an approach which could be used to describe novel pathogens in a wide range of different species.

## Author contributions

SRF conceived the study, performed all bioinformatic analyses and wrote the manuscript with contributions from all authors. SB, MC and JA provided the peafowl whole-genome sequencing data and provided analytical assistance. ALS provided conceptual and analytical assistance.

## Acknowledgements

SRF was supported by UKRI Biological Sciences Research Council (BBSRC) Institute Strategic Programme and Core Capability Grants to The Pirbright Institute (BBS/E/PI/230002B, BBS/E/PI/230002C and BBS/E/PI/23NB0003).

## References

Afonso, C. L., E. R. Tulman, Z. Lu, L. Zsak, D. L. Rock, and G. F. Kutish. 2001. “The Genome of Turkey Herpesvirus.” Journal of Virology 75 (2): 971–978.

Barbosa, Soraia, Carina Bittner, Roberto Arbore, et al. 2026. “Modulation of Avian Iridescence via Malanogenesis.” In bioRxiv. BioRxiv, January 27. 10.64898/2026.01.26.701322.

Bowman, Brian R., Matthew L. Baker, Frazer J. Rixon, Wah Chiu, and Florante A. Quiocho. 2003. “Structure of the Herpesvirus Major Capsid Protein.” The EMBO Journal 22 (4): 757–765.

Brito, Anderson F., Guy Baele, Kanika D. Nahata, Nathan D. Grubaugh, and John W. Pinney. 2021. “Intrahost Speciations and Host Switches Played an Important Role in the Evolution of Herpesviruses.” Virus Evolution 7 (1): veab025.

Brown, Andrew C., Lorraine P. Smith, Lydia Kgosana, Susan J. Baigent, Venugopal Nair, and Martin J. Allday. 2009. “Homodimerization of the Meq Viral Oncoprotein Is Necessary for Induction of T-Cell Lymphoma by Marek’s Disease Virus.” Journal of Virology 83 (21): 11142–11151.

Buchfink, Benjamin, Chao Xie, and Daniel H. Huson. 2015. “Fast and Sensitive Protein Alignment Using DIAMOND.” Nature Methods 12 (1): 59–60.

Casto, Amanda M., Pavitra Roychoudhury, Hong Xie, et al. 2020. “Large, Stable, Contemporary Interspecies Recombination Events in Circulating Human Herpes Simplex Viruses.” The Journal of Infectious Diseases 221 (8): 1271–1279.

Chen, X. B., P. J. Sondermeijer, and L. F. Velicer. 1992. “Identification of a Unique Marek’s Disease Virus Gene Which Encodes a 38-Kilodalton Phosphoprotein and Is Expressed in Both Lytically Infected Cells and Latently Infected Lymphoblastoid Tumor Cells.” Journal of Virology 66 (1): 85–94.

Dhama, Kuldeep, Naveen Kumar, Mani Saminathan, et al. 2017. “Duck Virus Enteritis (duck Plague) - a Comprehensive Update.” The Veterinary Quarterly 37 (1): 57–80.

Dotto-Maurel, Aurélie, Isabelle Arzul, Benjamin Morga, and Germain Chevignon. 2025. “Herpesviruses: Overview of Systematics, Genomic Complexity and Life Cycle.” Virology Journal 22 (1): 155.

Edgar, Robert C. 2004. “MUSCLE: Multiple Sequence Alignment with High Accuracy and High Throughput.” Nucleic Acids Research 32 (5): 1792–1797.

Fiddaman, Steven R., Evangelos A. Dimopoulos, Ophélie Lebrasseur, et al. 2023. “Ancient Chicken Remains Reveal the Origins of Virulence in Marek’s Disease Virus.” Science (New York, N.Y.) 382 (6676): 1276–1281.

Gimeno, I. M., R. L. Witter, H. D. Hunt, S. M. Reddy, L. F. Lee, and R. F. Silva. 2005. “The pp38 Gene of Marek’s Disease Virus (MDV) Is Necessary for Cytolytic Infection of B Cells and Maintenance of the Transformed State but Not for Cytolytic Infection of the Feather Follicle Epithelium and Horizontal Spread of MDV.” Journal of Virology 79 (7): 4545–4549.

Gowthaman, Vasudevan, Sachin Kumar, Monika Koul, et al. 2020. “Infectious Laryngotracheitis: Etiology, Epidemiology, Pathobiology, and Advances in Diagnosis and Control - a Comprehensive Review.” The Veterinary Quarterly 40 (1): 140–161.

Hyatt, Doug, Gwo-Liang Chen, Philip F. Locascio, Miriam L. Land, Frank W. Larimer, and Loren J. Hauser. 2010. “Prodigal: Prokaryotic Gene Recognition and Translation Initiation Site Identification.” BMC Bioinformatics 11 (1): 119.

Jones, D., L. Lee, J. L. Liu, H. J. Kung, and J. K. Tillotson. 1992. “Marek Disease Virus Encodes a Basic-Leucine Zipper Gene Resembling the Fos/jun Oncogenes That Is Highly Expressed in Lymphoblastoid Tumors.” Proceedings of the National Academy of Sciences of the United States of America 89 (9): 4042–4046.

Kaleta, Erhard F., and Douglas E. Docherty. 2008. “Avian Herpesviruses.” In Infectious Diseases of Wild Birds. Blackwell Publishing Professional.

Kingham, Brewster F., Vladimır Zelnık, Juraj Kopáček, Vladimır Majerčiak, Erik Ney, and Carl J. Schmidt. 2001. “The Genome of Herpesvirus of Turkeys: Comparative Analysis with Marek’s Disease Viruses.” The Journal of General Virology 82 (Pt 5): 1123–1135.

Li, Heng, and Richard Durbin. 2009. “Fast and Accurate Short Read Alignment with Burrows-Wheeler Transform.” Bioinformatics (Oxford, England) 25 (14): 1754–1760.

Li, Heng, Bob Handsaker, Alec Wysoker, et al. 2009. “The Sequence Alignment/Map Format and SAMtools.” Bioinformatics (Oxford, England) 25 (16): 2078–2079.

Lupiani, Blanca, Lucy F. Lee, Xiaoping Cui, et al. 2004. “Marek’s Disease Virus-Encoded Meq Gene Is Involved in Transformation of Lymphocytes but Is Dispensable for Replication.” Proceedings of the National Academy of Sciences of the United States of America 101 (32): 11815–11820.

Mo, Jongsuk, and Jongseo Mo. 2025. “Infectious Laryngotracheitis Virus and Avian Metapneumovirus: A Comprehensive Review.” Pathogens 14 (1): 55.

Nair, Venugopal. 2005. “Evolution of Marek’s Disease -- a Paradigm for Incessant Race between the Pathogen and the Host.” Veterinary Journal (London, England: 1997) 170 (2): 175–183.

Nair, Venugopal. 2018. “Spotlight on Avian Pathology: Marek’s Disease.” Avian Pathology: Journal of the W.V.P.A 47 (5): 440–442.

Nayfach, Stephen, Antonio Pedro Camargo, Frederik Schulz, Emiley Eloe-Fadrosh, Simon Roux, and Nikos C. Kyrpides. 2021. “CheckV Assesses the Quality and Completeness of Metagenome-Assembled Viral Genomes.” Nature Biotechnology 39 (5): 578–585.

Neerukonda, Sabari Nath, Upendra K. Katneni, Nirajan Bhandari, and Mark S. Parcells. 2019. “Transcriptional Analyses of Innate and Acquired Immune Patterning Elicited by Marek’s Disease Virus Vaccine Strains: Turkey Herpesvirus (HVT), Marek’s Disease Virus 2 (strain SB1), and Bivalent Vaccines (HVT/SB1 and HVT-LT/SB1).” Avian Diseases 63 (4): 670–680.

Parcells, M. S., S. F. Lin, R. L. Dienglewicz, et al. 2001. “Marek’s Disease Virus (MDV) Encodes an Interleukin-8 Homolog (vIL-8): Characterization of the vIL-8 Protein and a vIL-8 Deletion Mutant MDV.” Journal of Virology 75 (11): 5159–5173.

Prjibelski, Andrey, Dmitry Antipov, Dmitry Meleshko, Alla Lapidus, and Anton Korobeynikov. 2020. “Using SPAdes DE Novo Assembler.” Current Protocols in Bioinformatics 70 (1): e102.

Quinlan, Aaron R., and Ira M. Hall. 2010. “BEDTools: A Flexible Suite of Utilities for Comparing Genomic Features.” Bioinformatics (Oxford, England) 26 (6): 841–842.

Read, Andrew F., Susan J. Baigent, Claire Powers, et al. 2015. “Imperfect Vaccination Can Enhance the Transmission of Highly Virulent Pathogens.” PLoS Biology 13 (7): e1002198.

Reddy, Sanjay M., Yoshihiro Izumiya, and Blanca Lupiani. 2017. “Marek’s Disease Vaccines: Current Status, and Strategies for Improvement and Development of Vector Vaccines.” Veterinary Microbiology 206 (July): 113–120.

Seeber, Peter A., Benoît Quintard, Florian Sicks, Martin Dehnhard, Alex D. Greenwood, and Mathias Franz. 2018. “Environmental Stressors May Cause Equine Herpesvirus Reactivation in Captive Grévy’s Zebras (Equus Grevyi).” PeerJ 6 (e5422): e5422.

Seimon, T. A., D. McAloose, B. Raphael, et al. 2012. “A Novel Herpesvirus in 3 Species of Pheasants: Mountain Peacock Pheasant (Polyplectron Inopinatum), Malayan Peacock Pheasant (Polyplectron Malacense), and Congo Peafowl (Afropavo Congensis).” Veterinary Pathology 49 (3): 482–491.

Stamatakis, Alexandros. 2014. “RAxML Version 8: A Tool for Phylogenetic Analysis and Post-Analysis of Large Phylogenies.” Bioinformatics (Oxford, England) 30 (9): 1312–1313.

Szpara, Moriah L., Lance Parsons, and L. W. Enquist. 2010. “Sequence Variability in Clinical and Laboratory Isolates of Herpes Simplex Virus 1 Reveals New Mutations.” Journal of Virology 84 (10): 5303–5313.

Verweij, Marieke C., Daniëlle Horst, Bryan D. Griffin, et al. 2015. “Viral Inhibition of the Transporter Associated with Antigen Processing (TAP): A Striking Example of Functional Convergent Evolution.” PLoS Pathogens 11 (4): e1004743.

Wang, Jialing, Shuai Yuan, Dongjie Zhu, et al. 2018. “Structure of the Herpes Simplex Virus Type 2 C-Capsid with Capsid-Vertex-Specific Component.” Nature Communications 9 (1): 3668.

Widén, Frederik, Carlos G. das Neves, Francisco Ruiz-Fons, et al. 2012. “Herpesvirus Infections.” In Infectious Diseases of Wild Mammals and Birds in Europe. Preprint, Wiley, August 31. 10.1002/9781118342442.ch1.

